# Conformation-Dependent Donor Selectivity in the Xanthan Gum Glycosyltransferase GumK Revealed by AI-Based Docking

**DOI:** 10.64898/2026.04.10.717502

**Authors:** Davide Luciano, Tova Alenfalk, Gaston Courtade

## Abstract

The interdomain flexibility of GT-B fold glycosyltransferases regulates substrate binding and catalysis, yet the role of local structural variations in donor substrate specificity remains unclear. GumK, a GT70 enzyme from *Xanthomonas campestris*, exhibits local plasticity within its donor-binding pocket. We classify this plasticity into two conformational states, defined by the presence (closed state) or absence (open state) of a conserved hydrophobic interaction stabilizing the pocket. Using the AI-enhanced docking approach GNINA, we investigated the relationship between these states and substrate specificity by comparing UDP-glucuronate with five acidic and neutral ligand analogs. While docking scores showed limited discrimination among ligands, distance-based analysis between the sugar C6 atom and Lys307 revealed conformation-dependent trends. In the open state, negatively charged sugars preferentially interact with Lys307 via their carboxylate groups. Conversely, the closed state favors interactions with the pyrophosphate moiety. To provide orthogonal validation, we generated ensembles of GumK–substrate complexes using AI-based cofolding methods (AlphaFold3 and Boltz). While Boltz-2 affinities correctly ranked substrate specificity, only the GNINA-generated ensembles were consistent with physics-based simulations in capturing the conformational basis of selectivity. These results suggest that donor specificity arises from the interplay between ligand chemistry and binding-site plasticity rather than from a single rigid binding mode, and highlight complementary strengths of docking and cofolding approaches for studying flexible CAZymes.

## Introduction

Glycosyltransferases (GTs) catalyze the formation of glycosidic bonds and are essential for the biosynthesis of a wide range of glycoconjugates and polysaccharides. By transferring a monosaccharide unit from an activated donor, typically a nucleotide sugar, to a specific acceptor, these enzymes regulate the assembly of complex carbohydrate structures and their derivatives [1]. Due to their catalytic versatility, GTs are increasingly used as biocatalysts to produce glycans and functional biomaterials, and are also attractive targets for therapeutic intervention [2–4]. Despite their broad relevance, the molecular principles governing donor recognition and catalytic selectivity remain only partially understood.

GTs display a high degree of structural diversity and are classified into 139 families in the CAZy database based on sequence similarity [5]. These families adopt different structural folds, including GT-A, GT-B, GT-C, and other less common architectures [6], which reflect the wide range of catalytic strategies and substrate specificities within the superfamily.

Among the different architectures, the GT-B fold is one of the most abundant and conserved. GT-B enzymes are composed of two Rossmann-like domains connected by a flexible linker, with the catalytic site located at the interface between the two domains [7]. The N-terminal domain is typically involved in acceptor binding, whereas the C-terminal domain accommodates the donor substrate. In contrast to enzymes with a rigid and preorganized active site, GT-B proteins often rely on interdomain motions to regulate substrate binding and catalysis [8]. Despite the prevalence of GT-B enzymes, their conformational landscape and its role in determining substrate selectivity, particularly toward the donor substrate, remain incompletely understood.

The limited structural and mechanistic characterization of many GT-B enzymes poses a challenge for understanding their function and for exploiting them in biotechnological applications. In this context, the development of efficient strategies to rapidly screen donor–enzyme interactions would be highly valuable [9]. Computational approaches offer a promising starting point, providing atomistic insight into substrate recognition and enabling the exploration of multiple candidate substrates in a relatively high-throughput manner.

Among the less explored GT-B families, GT70 includes only one characterized member, GumK, a GT from *Xanthomonas campestris* involved in the biosynthesis of xanthan gum, a highly versatile microbial polysaccharide widely used in the food and materials industries [10, 11]. GumK catalyzes the transfer of glucuronic acid from UDP-glucuronate to a lipid-linked oligosaccharide intermediate, thus incorporating an acidic sugar moiety that contributes to the physicochemical properties of the final polymer [12]. The crystal structure of GumK (PDB ID: 2HY7)[13] revealed Asp157 as the catalytic base, consistent with an inverting reaction mechanism [14]. It has been experimentally shown that GumK binds UDP-glucuronate, but neither UDP-glucose nor UDP-galacturonic acid [15].

Understanding donor selectivity is of particular interest for both mechanistic and biotechnological reasons. While UDP-glucuronate is the native substrate, incorporating alternative UDP-sugars could enable the biosynthesis of modified xanthan derivatives with altered charge distribution and mechanical properties. The discrimination between acidic and neutral sugar moieties is expected to arise from specific electrostatic interactions within the donor-binding pocket, yet the residues responsible for this selectivity have not been conclusively identified [13, 15].

In a previous computational study [16], we investigated the conformational landscape of GumK and the conformational space of the monosaccharidic part of the donor substrate when the nucleotidic moiety is in the bound state at the isolated C-domain. We compared the ring orientation of UDP-glucuronate (the natural substrate) with that of the non-substrate UDP-glucose. We observed that the polarity of the C-domain’s binding site, together with its flexibility, may influence donor selectivity by favoring interaction between the carboxyl group of glucuronate and the key residue Lys307. This interaction may be important for both the catalytic mechanism and the substrate-binding pathway, providing an additional enthalpic contribution that is absent in UDP-glucose.

However, the computational strategy used to sample sugar conformations was not high-throughput, thereby limiting the utility of the approach for studying multiple donor substrate candidates or for screening mutants with potentially altered activity. To address this sampling limitation, we herein explore the use of docking as a complementary approach to investigate the interactions of various ligands with both full-length GumK and its isolated C-domain. We performed systematic docking calculations using GNINA [17], an AI-based scoring algorithm that has demonstrated improved performance over the standard AutoDock Vina scoring scheme. Our results show that the docking algorithm recovers pose populations similar to those generated by molecular dynamics (MD) simulations [16], enabling discrimination of substrates based on geometric descriptors. However, we found the outcome is highly sensitive to the conformation of the donor-binding site within the C-domain of GumK. Thanks to recent developments in AI-based co-folding methods that enable the prediction of protein–ligand complexes, we additionally employed AlphaFold3 and two versions of Boltz to generate multiple structural samples for each UDP-sugar ligand in complex with GumK. This comparison allows us to assess whether independent cofolding ensembles recover the same conformational states and substrate-dependent trends identified by GNINA, and to evaluate the utility of cofolding methods for saccharide–enzyme docking in the context of flexible CAZymes.Altogether, this study provides mechanistic insight into the dynamic determinants of donor recognition in a GT-B enzyme and offers a structural basis for the rational engineering of GumK toward alternative UDP-sugar substrates. Additionally, it provides insights into the strengths and limitations of cofolding methods and AI-based docking compared with physics-based molecular simulations for studying highly flexible carbohydrate–protein systems.

## Results and Discussion

### Conformational states of the donor-binding site

We focused our study on the donor-binding domain (C-domain) of GumK to isolate its binding properties from interactions with the acceptor domain. This choice also simplifies the highly dynamic two-domain enzyme into a relatively static one, thereby avoiding the need to sample multiple states with varying degrees of opening when studying the donor substrate’s conformation in the binding site [16].

The donor-binding domain of GumK shows a peculiar characteristic that is absent in the other xanthan gum GT-Bs, namely a flexible loop between residues 293–300, which shows high variability in terms of amino acid composition within the CAZy family of GumK (GT70) [5, 16]. This loop is not directly in contact with the donor substrate, as a hydrophobic interaction between Met231 and Leu301 positions it away from the donor-binding site. The crystal structures of GumK in the *apo* form (PDB: 2HY7) (Fig. 1A) and in complex with UDP (PDB: 2Q6V) (Fig. 1B) reveal that these hydrophobic residues adopt different conformational states. One conformation shows the side chain of Met231 interacting with that of Leu301 in the *apo* structure, whereas this interaction is absent in the UDP-bound complex. These residues are conserved within the GT70 family, and their conformational plasticity is partially attributable to the aforementioned flexible loop.

**Figure 1.**
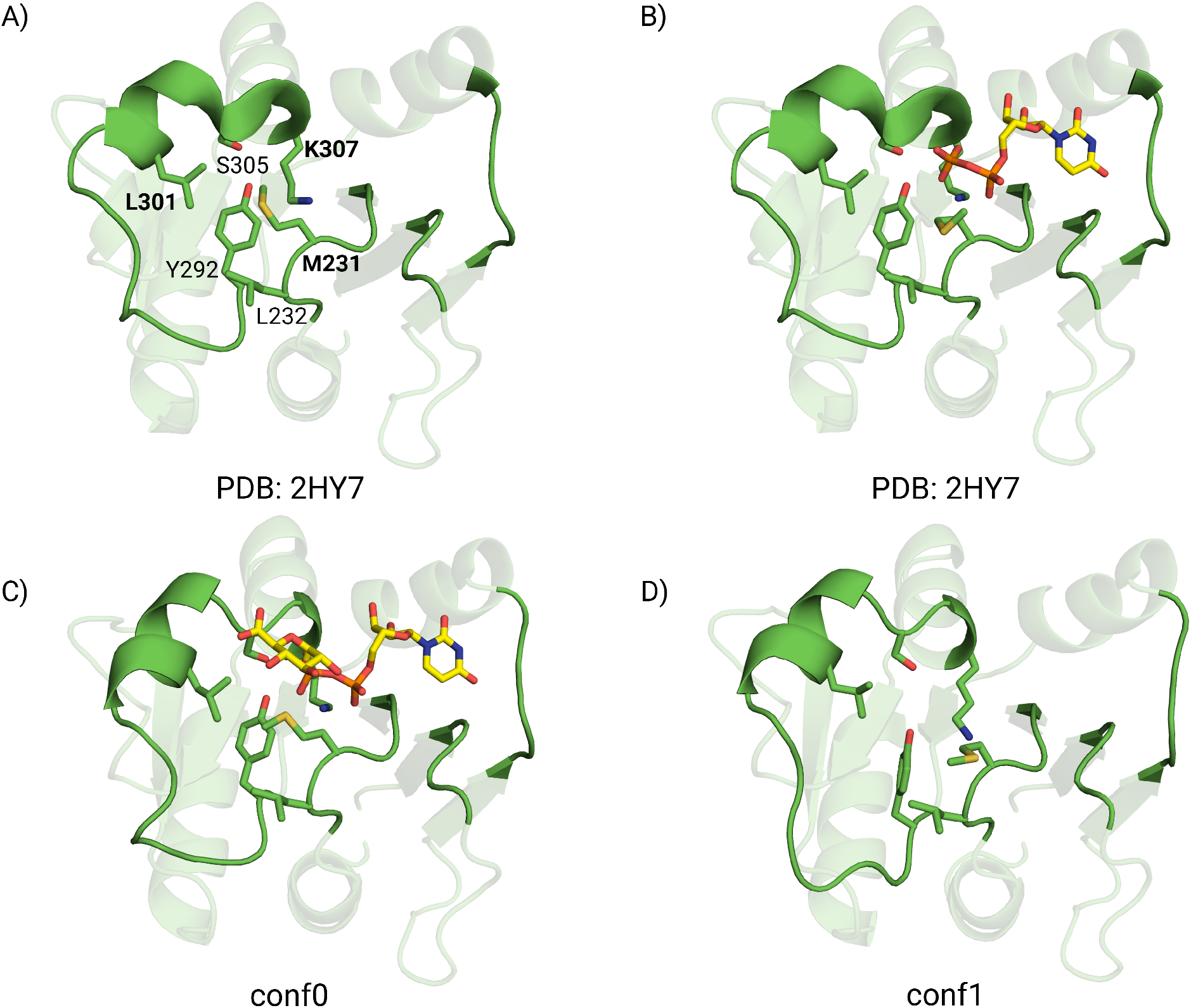
A) Conformation of the donor-binding domain taken from the *apo* crystal structure (PDB: 2HY7), with the key amino acids analyzed in this work labeled in bold. B) Conformation of the same domain in the crystal structure of GumK in complex with UDP (yellow) (PDB: 2Q6V). C) Boltz-2 prediction of the donor binding domain in complex with the donor substrate (UDP-GlcA in yellow). This conformation corresponds to conf0 in the main text. D) Donor-binding domain conformation sampled from a preparative simulation of the donor substrate in solution. This conformation corresponds to conf1 in the main text.

Patrick and colleagues used NMR spectroscopy to show that binding of the donor substrate is initially guided by the nucleobase, i.e., UDP, in a glycosyltransferase from *E. coli* (WaaG) that binds UDP-glucose [18]. Once this part is bound, the unconstrained sugar moiety retains conformational freedom, which is lost as the substrate approaches the catalytic pocket and finally binds.

Considering the conformational flexibility of both the binding site and the donor substrate, we evaluated the performance of GNINA in docking the native substrate and a series of UDP-sugars that are present in the metabolism of *Xanthomonas* as building blocks for several glycosylation reactions: UDP-glucose, UDP-galactose, UDP-galacturonate, UDP-*N*-acetylglucosamine, and UDP-*N*-acetylgalactosamine. Each ligand was docked onto two different states of the donor domain: one conformation obtained from the prediction of the C-domain in complex with full-length UDP-glucuronate using Boltz-2 (named conf0 in this work)[19] (Fig. 1C), and a second conformation extracted from a previously published [16] unbiased MD simulation of the isolated donor domain (named conf1 in this work) (Fig. 1D).

The conf1 corresponds to an open state of the hydrophobic interaction, confirming that this is a natural behavior of the donor-binding site and does not depend on the presence of the acceptor-binding domain, as suggested by the crystal structure [13]. In contrast, the Boltz2 cofolded complex, conf0, displays a closed state of the hydrophobic interaction. In this model, the sugar moiety does not directly interact with the binding domain, and binding is primarily mediated by the pyrophosphate group.

### GNINA docking recapitulates MD-derived pose populations

For each ligand, a GNINA docking approach was employed to obtain a statistically meaningful representation of the accessible binding poses. We ran 31 replicas per ligand, with each replica generating 20 poses, yielding a total of 31 × 20 docked poses per ligand. We plotted the distributions of the median GNINA CNN score (convolutional neural network score) computed for each replica, i.e., over 20 poses (Fig. 2A and B). We filtered out poses in which the nucleobase was not in the crystallographic binding mode, based on RMSD relative to the crystal structure (PDB 2Q6V). For the remaining poses, we computed a bootstrapped distribution of the distance between the C6 atom of the ligand and the N*η* atom of the key residue Lys307 (*d*_*K*_) (Fig. 2C and D).

**Figure 2.**
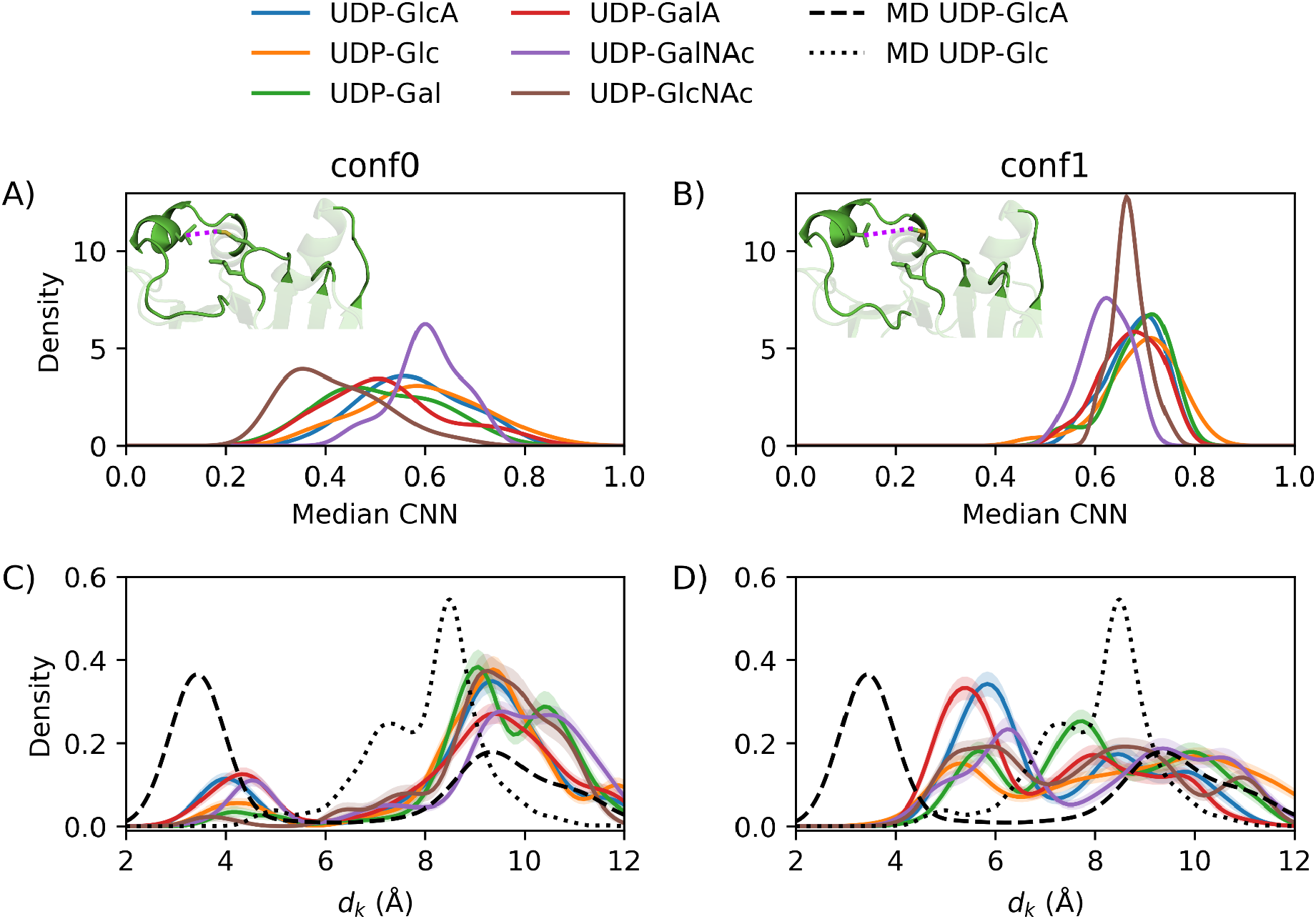
A) Distribution of the median GNINA CNN score across docking replicas using two different conformations of the donor binding site. conf0 has a short distance between Met231 and Leu301, as shown by the structure in panel A. conf1 has a longer distance between the two residues, as shown by the dashed line in the structure in panel B. In the left column, results using conf0 as the receptor; in the right column, results using conf1 as the receptor. Each replica generates 20 poses, and the median score is calculated over these poses using conf0 as the receptor. The resulting values are compared across ligands, with UDP-GlcA used as a reference. B) Like A, but using conf1 as the receptor in the docking calculations. C) Bootstrapped distribution (4,000 resampling cycles) of the distance between the C6 atom of the tested ligands and the *ζ*-nitrogen of Lys307 (*d*_*K*_ in Å), calculated over all conformations in which the uridine moiety is correctly docked based on the RMSD respect to the crystallographic mode. The shaded region represents the standard deviation obtained from the bootstrap procedure. D) Like C, but using conf1 as the receptor. The dotted and dashed curves correspond to the weighted distributions of the same variable (*d*_*K*_) sampled during Hamiltonian replica exchange simulations of UDP-GlcA and UDP-Glc with the uridine moiety constrained, as reported in our previous work [16].

Inspection of the distributions of median CNN scores across ligands reveals no apparent difference. The only distribution that deviates from the native substrate (UDP-GlcA) is UDP-GlcNAc in the conf0 state of the binding site. Assuming that the donor domain specifically binds UDP-GlcA, then the score alone cannot be used to discriminate between substrates, as it fails to distinguish between a good binder and a non-binder.

However, when analyzing the distribution of the reference distance (*d*_*K*_), clear differences emerge. In the closed state of the binding site (conf0; Fig. 2C), two peaks at approximately 4 Å are observed for the acidic ligands UDP-GlcA and UDP-GalA, corresponding to a direct interaction between the carboxyl group and Lys307. In this C-domain conformation, the neutral substrates display a lower population in this region, while their main peak is centered above 8 Å. At these distances, Lys307 interacts primarily with one or both phosphate groups of the pyrophosphate moiety.

In the open conformation (conf1), the distributions of the two acidic ligands change substantially, and their dominant peaks become associated with direct interactions between the carboxyl group and Lys307 (Fig. 2D). In contrast, the neutral substrates exhibit broader distributions spanning a wider range of distances *d*_*K*_. This behavior likely reflects the increased conformational space available for reorienting the sugar moiety in conf1 relative to conf0.

Filtering the conformations based on the docked orientation of the nucleobase in the crystal structure has a similar effect to constraining the nucleobase’s RMSD to the reference conformation, a strategy previously used to study the interaction using MD simulations [16]. The conformations generated with GNINA are in agreement with the states sampled in MD simulations (Fig. 2C and D). The region between 8 and 10 Å represents interactions with the phosphate groups, whereas in the region between 2 and 4 Å the C6 atom of the monosaccharide is located close to Lys307. The GNINA-generated distributions do not completely overlap with those generated by MD because, in the latter, the flexible side chain of Lys307 can rearrange to interact optimally with the carboxyl group of UDP-GlcA, maintaining conformation, whereas in the former, the side chains are rigid. Despite this difference, the bimodal nature of the distributions is recapitulated.

In MD simulations, the sampled function corresponds to a biased potential energy surface (using Hamiltonian Replica Exchange [16]) defined by a force field that models the physical interactions of the system. In contrast, GNINA employs a CNN-based scoring function in which atomic positions within the docking box are associated with the likelihood of observing that distribution in experimental crystal structures. Specifically, the CNN predicts both a pose score (a confidence score between 0 and 1 that a pose has a low RMSD relative to the experimental structure) and binding affinity in pK units [20]. The scoring function, therefore, contains no explicit physical energy terms.

Although the CNN score could not discriminate among substrates, it provided useful insights into binding-site selectivity. As the protein transitions from the open (conf1) to closed (conf0) conformation, the median CNN score distributions become less overlapping (Fig. 2A–B). However, due to system flexibility and multiple local minima, poses with an improperly bound nucleobase can still achieve high CNN scores and vice versa, since the nucleobase’s binding contribution can be partially compensated by the sugar–pyrophosphate moiety. By introducing an RMSD filter and a suitable binding-mode descriptor, GNINA nonetheless recovered a conformation distribution resembling that of a physical potential energy surface. This likely reflects the CNN’s complex representation of the chemical environment, though extracting interpretable information for a specific docked system remains challenging [21].

Carbohydrates are cyclic molecules whose ring structures impose important conformational constraints. A potential limitation of scoring functions that are not explicitly physics-based is that they may generate highly distorted ligand conformations or fail to preserve the correct stereochemistry. Consequently, structural filtering is often required to identify and remove physically unrealistic poses [22]. However, for GNINA, we did not observe significant distortions, suggesting that the model is reasonably well-suited to these types of molecules. The limited quantitative discrimination observed for the CNN score may be related to the nature of the training dataset. Structural information for GT-B glycosyltransferases remains relatively scarce, and GumK is currently the only member of the CAZy GT70 family for which an experimental structure is available. As a result, binding modes and structural features specific to this enzyme family are likely underrepresented in the data used to train the scoring model, potentially affecting its performance on these systems [23].

Despite these limitations, we demonstrated that prior knowledge from more computationally demanding physics-based simulations, combined with a discriminating variable to characterize the generated conformations, yielded an AI-based docking approach that provides meaningful insights into the behavior of these systems while retaining the advantages of high-throughput screening.

### Cofolding predicts substrate affinity but not conformational specificity

To provide orthogonal validation of the docking-based analysis, we generated ensembles of GumK–substrate complexes using AlphaFold3 [24] and Boltz-2 [19] by producing 200 diffusion samples per UDP-sugar substrate. This approach was motivated by the hypothesis that a sufficiently large number of samples could capture alternative interdomain conformations, yielding a receptor ensemble implicitly coupled to each substrate.

Inspection of the predicted interdomain geometries (Fig. 3) revealed that cofolding samples predominantly converge on narrow distance distributions between the C6 atom of the monosaccharides and Lys307, consistent with non-binding conformations. This behavior likely reflects the tendency of diffusion-based structure prediction models to reproduce conformations seen in the experimental structures [25]. As a consequence, the substrate-dependent conformational trends identified by GNINA (Fig. 2) were not recovered from the cofolding ensembles. These methods are also prone to well-documented stereochemical errors [25, 26], which we also observed in our predicted ensembles of AlphaFold3 and Boltz-2. However, as we did not observe chirality violations in Boltz-1, we also included ensembles of 100 diffusion samples generated by this method. Yet, its *d*_*K*_ distributions remained nearly identical across all six UDP-sugars (Fig. 3), confirming that stereochemical fidelity alone cannot recover substrate-dependent conformational discrimination.

**Figure 3.**
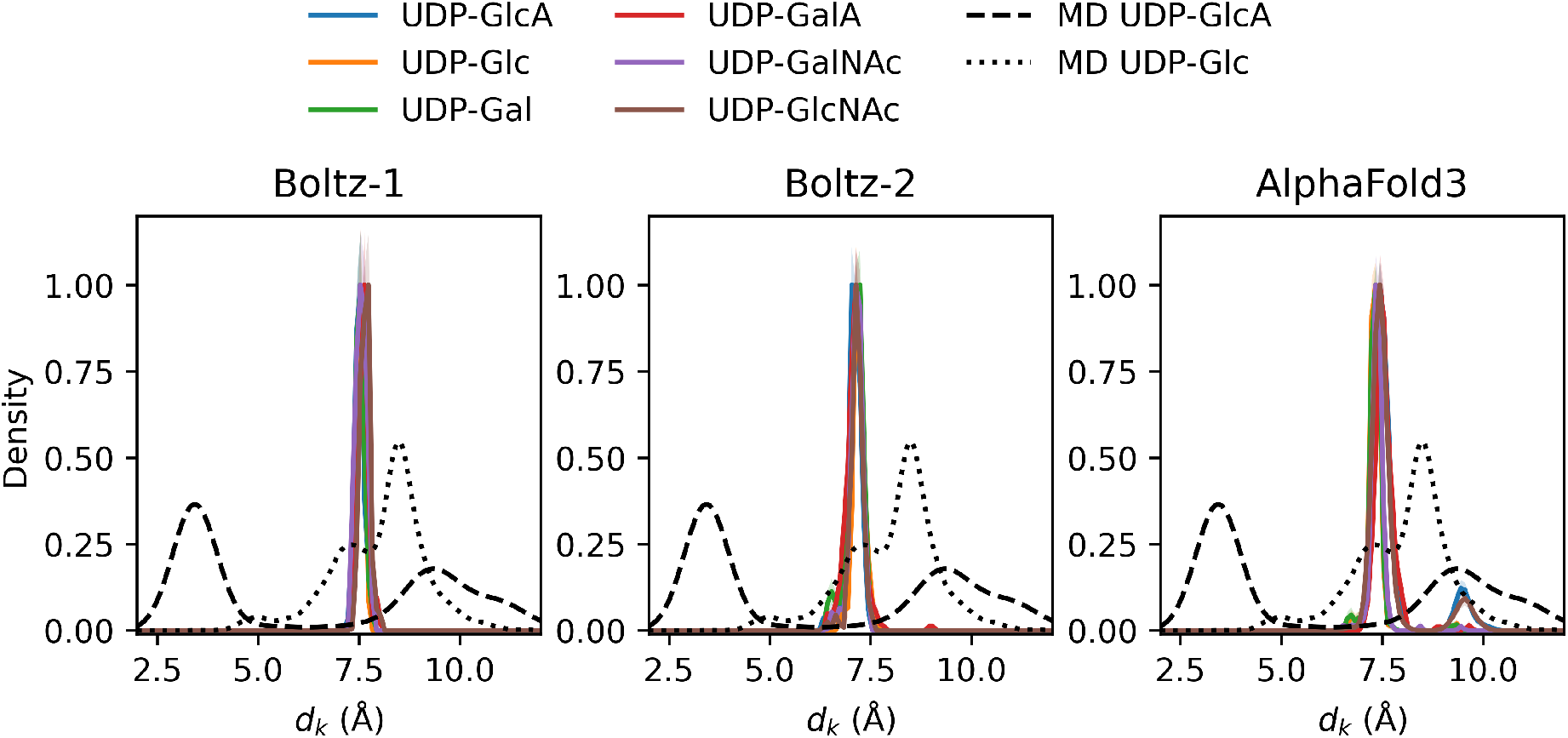
Bootstrapped distribution (4,000 resampling cycles) of the distance between the C6 atom of the tested ligands and the *ζ*-nitrogen of Lys307 (*d*_*K*_ in Å), calculated over all conformations in the ensembles generated by Boltz-1 (N=100), Boltz-2 (N=200), and AlphaFold3 (N=200). The shaded region represents the standard deviation obtained from the bootstrap procedure. The dotted and dashed curves correspond to the weighted distributions of the same variable (*d*_*K*_) sampled during Hamiltonian replica exchange simulations of UDP-GlcA and UDP-Glc with the uridine moiety constrained, as reported in our previous work [16].

Nevertheless, Boltz-2 affinity scores (Table 1) correctly ranked the substrates, assigning the highest predicted affinity to the native substrate UDP-GlcA, closely followed by UDP-GalA, and lower affinities to the neutral analogs. This result is consistent with the hypothesis that negatively charged ligands are preferentially recognized by GumK and aligns with the enthalpic contribution attributed to the Lys307–carboxylate interaction identified in the docking and MD analyses. However, the mechanistic basis for this ranking cannot be directly extracted from the cofolding output, as the score is not decomposable into conformation-dependent contributions, in contrast to the geometric descriptor *d*_*K*_.

**Table 1.** Affinity scores calculated by Boltz-2 for each of the ligands. Scores can be approximately interpreted as a *log*_10_(*IC*_50_), derived from an *IC*_50_ measured in *µ*M [19]

| Ligand | Boltz-2 Affinity |
| --- | --- |
| GlcA | −0.6550 |
| GalA | −0.5980 |
| Glc | −0.2537 |
| Gal | −0.2512 |
| GalNAc | −0.1721 |
| GlcNAc | −0.1349 |

Taken together, these results suggest that GNINA docking and AI-based cofolding provide complementary rather than equivalent information. Boltz-2 offers a rapid route to affinity ranking, but cofolding methods cannot capture the conformational plasticity of flexible binding sites, a critical feature for predicting donor specificity in GT-B glycosyltransferases. Additionally, carbohydrates are a challenging class of molecules for AI-based models due to their high degree of chirality, which may not always be accurately captured by deep learning approaches. Nevertheless, cofolding models can be highly valuable for generating initial protein–ligand conformations that can subsequently be refined and analyzed using docking calculations or molecular simulations.

### GNINA docking on full-length GumK shows the effect of the N-domain

The docking experiments on the isolated C-domain enabled us to isolate the effect of the binding-site conformation on donor-substrate binding and potential substrate selectivity. However, the binding site of the isolated C-domain is largely solvent-exposed, allowing a greater conformational freedom for the sugar moiety. In contrast, the presence of the N-domain in the closed conformation may contribute to substrate selectivity by further constraining the orientation of the ligand. To test this hypothesis, we generated full-length GumK–ligand complexes by cofolding UDP-GlcA and UDP-GalA with Boltz-1 and subsequently performed docking calculations to evaluate how the N-domain affects the GNINA-generated distributions (Fig. 4).

**Figure 4.**
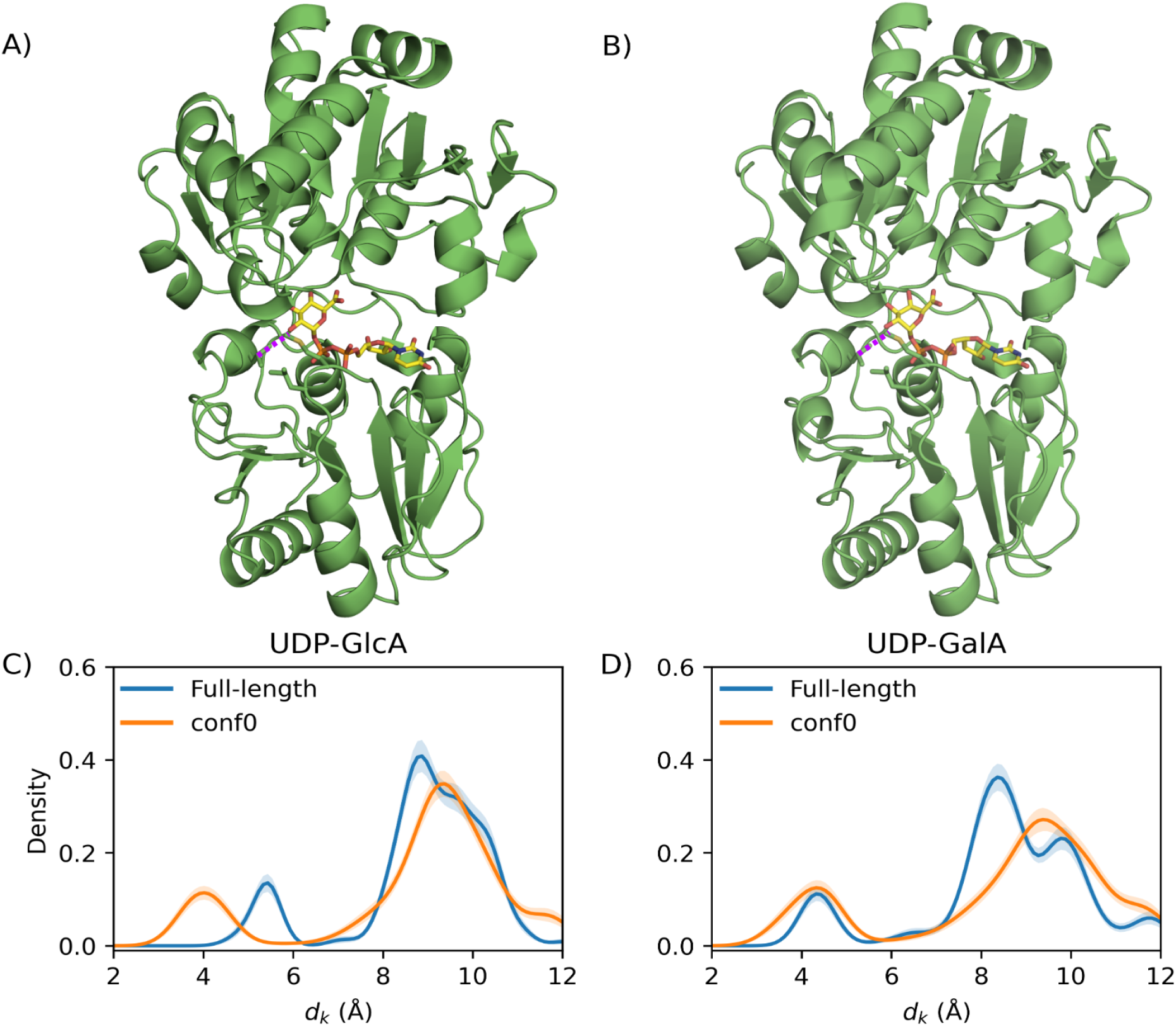
A) Boltz-1 cofolded complex of UDP-GlcA bound to full-length GumK. The distance between Met298 and Leu301 is highlighted with a dashed line, indicating that the C-domain adopts a closed conformation, similar to that observed in conf0. B) Boltz-1 cofolded complex of UDP-GalA bound to full-length GumK. The same distance between the key hydrophobic residues is highlighted to assess the conformational state of the C-domain. In this case the ring of the sugar moiety was predicted in a distorted chain representation. C) Comparison of the bootstrapped *d*_*k*_ distributions obtained from the docked conformations of UDP-GlcA with full-length GumK and conf0 for the isolated C-domain (Fig. 2C). D) Comparison of the bootstrapped *d*_*k*_ distributions obtained from the docked conformations of UDP-GalA with full-length GumK and conf0 (Fig. 2C). The shaded region represents the standard deviation obtained from the bootstrap procedure.

The cofolded structures adopt a closed-state conformation, in which the pyrophosphate region of the ligand interacts with Lys307. Based on the Met231–Leu301 distance, the resulting structures correspond to the conf0 state of the C-domain.

Redocking with GNINA, using the same protocol as for the isolated C-domain, revealed subtle but noticeable differences in the distance distributions. In particular, the interaction between the carboxyl group and Lys307 is less populated in the case of UDP-GlcA (Fig. 4A), and both ligands display a dominant peak around 8 Å, consistent with interactions between Lys307 and the pyrophosphate moiety. This shift in the distribution may be explained by the additional contacts provided by the N-domain, which, as expected, further restricts the orientation of the ligand within the binding site.

### Ensemble docking considerations

Although the lack of receptor flexibility represents an intrinsic limitation of the docking approach, it may also provide a conceptual advantage. By constraining the protein conformation, docking enables a more controlled comparison of how predefined binding-site states influence the spatial distribution of the docked ligands, thereby facilitating the dissection of conformation-dependent specificity.

The docking study suggests that acidic ligands contribute an additional enthalpic component to binding through interactions between the carboxylate and Lys307. This interaction may influence the binding pathway by properly orienting the substrate for catalysis. The flexibility of the donor-binding site may guide this interaction through two steps: in a more open state, the sugar moiety can interact with Lys307 via its carboxyl group, as there is sufficient space to accommodate the sugar ring in the proper orientation. When the Met-Leu loop adopts a closed conformation, the interaction with the carboxyl group is reduced, whereas interactions with the pyrophosphate group become more favored, allowing the sugar ring to approach the catalytic residue in the acceptor-binding domain (Fig. 5).

**Figure 5.**
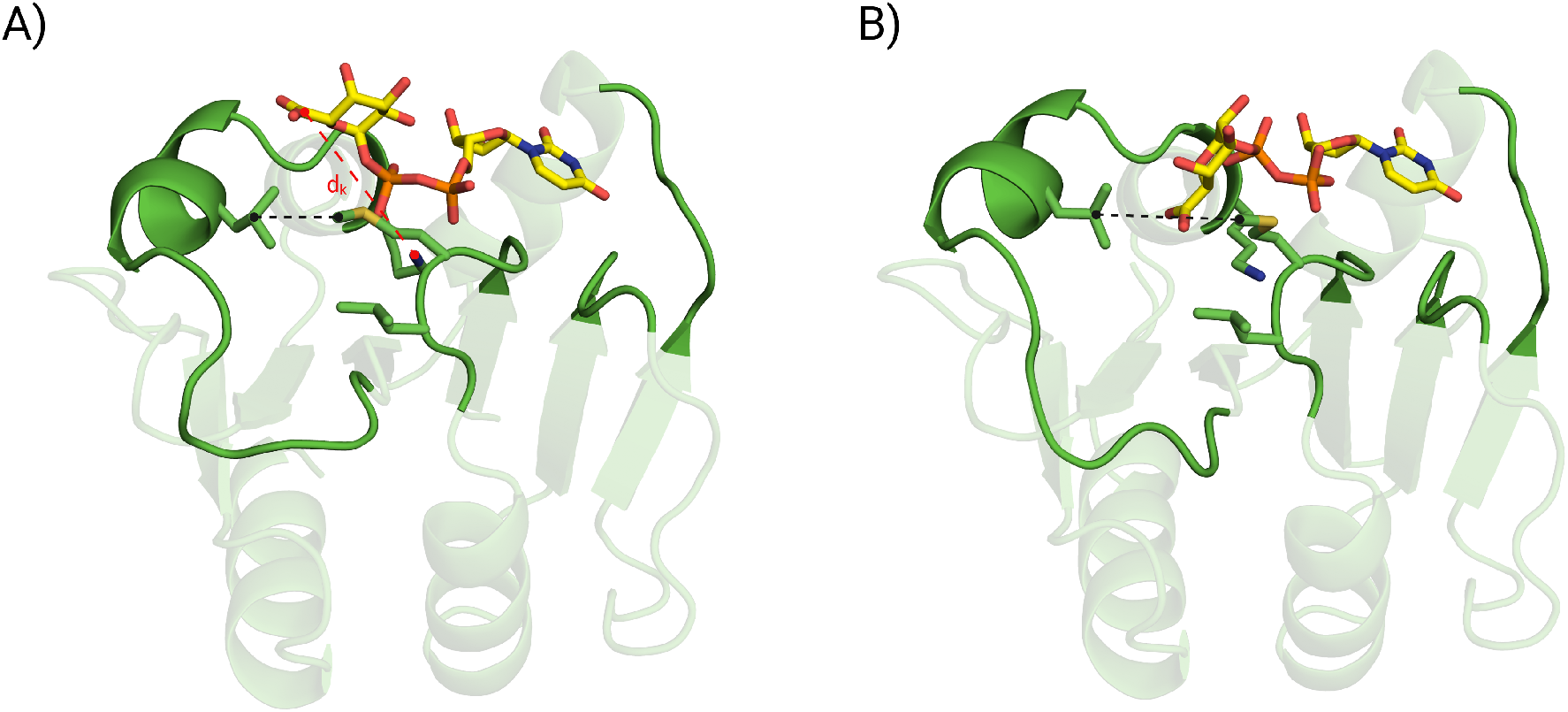
A) Representative docked conformation of UDP-glucuronate interacting with conf0 through the pyrophosphate groups. The dashed red line shows the reference distance, *d*_*k*_, used to describe the binding modes. The black dashed line highlights the degree of opening of the binding site. B) Representative docked conformation of UDP-glucuronate interacting with conf1 through the carboxyl group.

This two-state behavior can be formalized within an ensemble-docking framework in which each GNINA-derived distribution represents the residual conformational freedom of the substrate conditioned on a specific protein conformation. The equilibrium substrate distribution can then be reconstructed as a weighted average of these conditional distributions, with weights given by the Boltzmann populations of the corresponding protein states, which can be estimated from molecular dynamics simulations. For example, conformations such as conf0 and conf1, as well as other conformational states of the full-length protein, would contribute proportionally to their equilibrium populations.

In practice, however, such an ensemble-docking strategy is non-trivial. The positioning of the uridine moiety within its binding pocket is not independent of the geometry of the surrounding donor-binding site, implying a structural coupling between local and global conformational features. Consequently, sampling only selected regions of the binding site without accounting for their collective organization is inherently challenging.

## Conclusion

In this study, we explored the potential of GNINA to investigate donor recognition in a system characterized by substantial conformational flexibility at both the binding site and the substrate level. GumK presents a challenging case for rigid docking approaches, as the donor-binding domain can adopt multiple conformations, and the UDP-sugar retains residual flexibility even after nucleobase anchoring.

To account for this structural variability, we represented the conformational heterogeneity of the donor-binding site through two structurally defined states. These were operationally defined based on the presence or absence of a conserved hydrophobic interaction that stabilizes the pocket: an open state, in which this interaction is disrupted, and a closed state, in which it is maintained. By docking multiple UDP-sugars to these two predefined conformations, we aimed to assess whether GNINA could capture substrate-dependent trends across distinct structural contexts.

While the median CNN score does not clearly discriminate between substrates, analysis of geometric descriptors suggests that substrate behavior depends on the binding pocket’s conformational state. In the closed state (conf0; Fig. 5A), the contacts between pyrophosphate and Lys307 represent the main population. In contrast, in the open state (conf1; Fig. 5B), acidic sugars more frequently adopt conformations compatible with an interaction between the carboxyl group and Lys307.

A similar picture emerges for the full-length protein, where the constraints on the conformational freedom of the sugar ligand arise not only from the architecture of the binding site within a single domain, but also from the presence of the second domain. This effect can be qualitatively appreciated from the changes observed in the distributions of the geometric descriptor used to characterize the sampled binding modes.

These observations are consistent with a model in which donor recognition may arise from the interplay between substrate chemistry and the binding site’s conformation, rather than from a single static binding mode. Although GNINA does not explicitly sample backbone motions, the two-conformation strategy and the full-length protein experiment adopted here suggest that docking can still capture meaningful trends when different structural states are explicitly considered.

Furthermore, we compared the AI-based docking approach with the cofolding methods implemented in AlphaFold3 and Boltz. In contrast to docking algorithms, where AI is primarily used to score conformations generated by a sampling procedure, cofolding methods rely on the AI model itself to directly sample the conformational space of the protein–ligand complex. In our case, these models appear to be biased toward a very narrow binding mode, likely reflecting the limited amount of experimental structural information available for such systems. In addition, they occasionally fail to preserve the correct stereochemistry of the ligands.

Despite not being explicitly physics-based, GNINA is surprisingly capable of generating an ensemble of binding modes that resembles those sampled during molecular dynamics simulations. This highlights the value of combining AI models with knowledge obtained from more computationally demanding methods and suggests that there is still considerable room for improvement in current cofolding approaches. Importantly, our results show that simple affinity scores alone are insufficient to discriminate between favorable and unfavorable ligands. Instead, meaningful interpretation requires analysis of the generated conformations using appropriate geometric descriptors. Importantly, due to its computational efficiency, the GNINA-based approach used in this work could, in principle, be extended to rapid screening of mutants, provided that the conformational ensemble of the donor-binding site is adequately represented. In this context, docking would not replace detailed molecular simulations, but could serve as an initial filter to identify variants whose binding-site conformations preserve or alter key substrate-dependent interactions.

Overall, this work suggests that AI-based docking, combined with an explicit representation of the binding site’s open and closed states, may provide a practical, exploratory framework for generating testable hypotheses about donor selectivity in flexible GT-B enzymes.

## Methods

### GNINA

The first conformation of the C-domain (conf0) was predicted using Boltz-2 [19], which takes as input only the sequence of the isolated domain and the CCD definition of UDP-GlcA (see below). The top-ranked model was selected and used as a receptor for docking. conf1 was instead extracted from a preparative MD simulation of the C-domain in solution.

Each ligand was docked using GNINA (version 1.3) [17]. The docking search space was defined by an autobox centered on residues surrounding the donor-binding site (228–231, 253– 254, 271–275, 278, 292, 301, 304–308, and 310), with automatic box extension enabled. The exhaustiveness parameter was set to 16, and the minimum RMSD between poses during the Monte Carlo sampling was set to 1 Å to filter redundant conformations. For each ligand, 31 independent docking runs were performed, each initialized with a different random seed and generating 20 poses per replica.

Since both receptor conformations (conf0 and conf1) were structurally aligned to the C-domain domain of the crystal structure (PDB ID: 2Q6V), the RMSD of the uridine moiety with respect to the crystallographic reference was used as a geometric filter. Among the poses generated for each replica, only those with a CNN score above the median and with the uridine moiety correctly positioned within the binding site were retained for further analysis.

### Cofolding methods

AlphaFold3 (v3.0.1) [24], Boltz-1 (v1.0.0) [27], and Boltz-2 (v2.2.1) [19] were used to predict cofolded structures of conf0 (see above) with each of the six UDP-sugar ligands, identified by their CCD codes (UGA: UDP-GlcA, UGB: UDP-GalA, UD1: UDP-GlcNAc, UD2: UDP-GalNAc, UPG: UDP-Glc, GDU: UDP-Gal). To generate conformational ensembles, the number of diffusion samples was set to 200 for AlphaFold3 and Boltz-2, and to 100 for Boltz-1 due to memory limitations. The resulting samples were aligned using PyMOL [28].

### Full-legth Gumk docking

The full-length GumK complexes with UDP-GlcA and UDP-GalA were predicted using Boltz-1 (v1.0.0), employing the CCD codes UGA and UGB for the respective ligands. Three diffusion samples were generated for each complex, although only the top-ranked model was selected for the docking analysis. After removing the cofolded ligand, GNINA docking calculations were performed using the same protocol adopted for the isolated C-domain. The resulting poses were filtered based on the uridine binding mode by calculating the RMSD of the uridine moiety relative to the crystallographic reference structure (PDB ID: 2Q6V). Only poses reproducing the reference uridine conformation were retained for subsequent analysis.

### Distance distribution analysis

For the filtered poses generated by the GNINA workflow and the conformational ensembles produced by the cofolding methods, the distance between the C6 atom of the ligand and the *ζ*-nitrogen of Lys307 was measured. Kernel density estimation (KDE) was then computed to characterize the distribution of this distance. To assess variability in the sampled conformations, a bootstrapped KDE was computed using 4000 resampling cycles, and the standard deviation was estimated from the resulting distribution.

## Data Availability Statement

Data and scripts used for running and analyzing simulations, and for generating the figures, are available from https://github.com/gcourtade/papers/tree/master/2026/GumK_GNINA

## Acknowledgements

G.C. gratefully acknowledges funding by the Novo Nordisk Foundation (grant number NNF22OC0073963).

